# Perturbations both trigger and delay seizures due to generic properties of slow-fast relaxation oscillators

**DOI:** 10.1101/2020.12.02.407965

**Authors:** Alberto Pérez-Cervera, Jaroslav Hlinka

## Abstract

The mechanism underlying the emergence of seizures is one of the most important unresolved issues in epilepsy research. In this paper, we study how perturbations, exogenous of endogenous, may promote or delay seizure emergence. To this aim, due to the increasingly adopted view of epileptic dynamics in terms of slow-fast systems, we perform a theoretical analysis of the phase response of a generic relaxation oscillator. As relaxation oscillators are effectively bistable systems at the fast time scale, it is intuitive that perturbations of the non-seizing state with a suitable direction and amplitude may cause an immediate transition to seizure. By contrast, and perhaps less intuitively, smaller amplitude perturbations have been found to delay the spontaneous seizure initiation. By studying the isochrons of relaxation oscillators, we show that this is a generic phenomenon, with the size of such delay depending on the slow flow component. Therefore, depending on perturbation amplitudes, frequency and timing, a train of perturbations causes an occurrence increase, decrease or complete suppression of seizures. This dependence lends itself to analysis and mechanistic understanding through methods outlined in this paper. We illustrate this methodology by computing the isochrons, phase response curves and the response to perturbations in several epileptic models possessing different slow vector fields. While our theoretical results are applicable to any planar relaxation oscillator, in the motivating context of epilepsy they elucidate mechanisms of triggering and abating seizures, thus suggesting stimulation strategies with effects ranging from mere delaying to full suppression of seizures.

**Author summary:** Despite its simplicity, the modelling of epileptic dynamics as a slow-fast transition between low and high activity states mediated by some slow feedback variable is a relatively novel albeit fruitful approach. This study is the first, to our knowledge, characterizing the response of such slow-fast models of epileptic brain to perturbations by computing its isochrons. Besides its numerical computation, we theoretically determine which factors shape the geometry of isochrons for planar slow-fast oscillators. As a consequence, we introduce a theoretical approach providing a clear understanding of the response of perturbations of slow-fast oscillators. Within the epilepsy context, this elucidates the origin of the contradictory role of interictal epileptiform discharges in the transition to seizure, manifested by both pro-convulsive and anti-convulsive effect depending on the amplitude, frequency and timing. More generally, this paper provides theoretical framework highlighting the role of the of the slow flow component on the response to perturbations in relaxation oscillators, pointing to the general phenomena in such slow-fast oscillators ubiquitous in biological systems.

## Introduction

The dynamics underlying complex processes usually involve many different time scales due to multiple constituents and their diverse interactions. When modelling such systems, the general distinction of at least two time-scales (fast and slow) is a useful and common conceptualization. Many examples of slow-fast dynamics can be found in cell modelling, ecosystems, climate or chemical reactions [1–4] and more recently in epilepsy [5], of particular interest for this paper.

Epilepsy is a chronic neurological disorder characterized by a marked shift in brain dynamics caused by an excessively active and synchronized neuronal population [6, 7]. Although several dynamical pathways have been proposed to explain the transition to seizure [8–10], in general, epilepsy is modelled as a system having two stable states: one corresponding to the low activity state and the other corresponding to high activity (that is to seizure) [11]. Besides external perturbations or noise, transitions between these two stable states can also be modelled considering the existence of a parameter evolving on some slow time scale. Whereas on the fast time scale the system can be seen as a bistable system, the variations of the slow parameter lead to bifurcations providing transitions between states [12].

During the last decade, there has been an increasing number of models approaching epilepsy through slow-fast time scales [13–16]. Recently, the slow-fast dynamics has been proposed to explain the role of the interictal epileptiform discharges (IEDs) in the generation of seizures [17]. The IEDs can be thought of as endogenous inputs affecting the target tissue. However, the effect of IEDs on the tissue activity is quite controversial: where some studies show that IEDs can prevent seizures [18, 19], other studies claim their seizure facilitating role [20, 21]. In the above mentioned work [17], the amplitude and frequency dependence of the effect of perturbations in a simple epilepsy model switching between seizure and non-seizure states due to a slow feedback variable, provided a conceptual reconciliation of the pro-convulsive and anti-convulsive effect of IEDs.

In this paper we elucidate this phenomena in detail and provide theoretical foundations of this apparent perturbation effect paradox by studying the phase response of a generic relaxation oscillator. We perform this theoretical approach by means of the phase reduction [22]. In addition to simplifying the dynamics, the usage of phase reduction techniques allows the computation of its *isochrons* and *phase response curves* (PRCs), which clarify the dependence of the effect of perturbations of the oscillator on the perturbation timing, and also allows the study of possible synchronization regimes [23]. By studying a generic slow-fast system displaying relaxation oscillations we show, analytically, how the slow component of the vector field shapes their isochrons and PRCs, thus ultimately determining its response to perturbations. Therefore, our results, clarify the multifaceted effect of IEDs in epilepsy, and can be straightforwardly applied to understand the temporal dependency of perturbations over any model belonging to the wide family of models relying on slow-fast dynamics.

The paper is structured as follows. First, we present a general introduction to relaxation oscillators introducing the basic notation which will be used throughout the paper. Then, we describe the phenomenological epilepsy model and show how, through its phase analysis, we can unveil the mechanism integrating the contradictory role of IEDs in epilepsy. Next, we show, via a complete theoretical analysis, which factors determine the geometry of isochrons of planar relaxation oscillators and study the response of perturbations of relaxation oscillators. We support our theoretical findings studying the response of perturbations for a different reduced epileptor model and discuss our results in the context of epilepsy. We conclude the paper by explaining the computational techniques in the Methods section.

## Results

### Basic introduction to relaxation oscillations

The main aim of this Section is to introduce the reader to the basics of slow-fast systems and in particular to relaxation oscillations. For further details we refer the reader to [24–27]. We will consider systems in this form

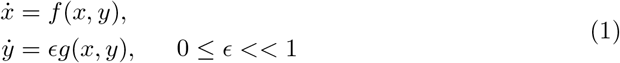

the flow of which will be denoted as *ϕ_t_*(*x, y*). Notice that indicates the derivative with respect to the time, *t*. As 0 ≤ *ϵ* << 1, the variables *x* and *y* evolve on different time-scales, namely the fast time, *t*, and the slow time *τ* = *ϵt*. Next, we use this distinction between time-scales to illustrate how a system in the form (1) with the extra assumption of *f*(*x, y*) = 0 being a cubic manifold, generates a limit cycle (denoted as Γ_*ϵ*_) showing relaxation oscillations [28] (see also Fig. 1). Consider a point *p* = (*x, y*). First, since *ϵ* << 1, we can take the limit *ϵ* → 0 and approximate the dynamics of system (1) by the *layer* dynamics

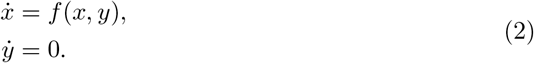

The trajectory of *p* will initially (approximately) follow the layer dynamics in (2) so it will quickly converge to its set of equilibrium points, defined as the *slow manifold* 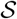

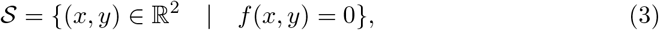

which in the limit *ϵ* → 0 corresponds to the nullcline (*ẋ* = 0) of the fast variable. As we considered the slow manifold 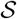 in (3) to be cubic, that is S-shaped, it will have two fold points (given by *∂_x_f*(*x, y*) = 0), which we denote as 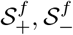 respectively, separating the repelling and attracting branches, denoted as 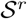 and 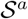, respectively

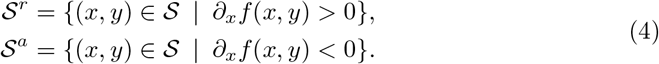

Note that the attracting part of the slow manifold 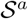 in fact consists of a top and bottom branch 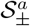. Once the system has approached the slow manifold, its dynamics are given by the slow variable

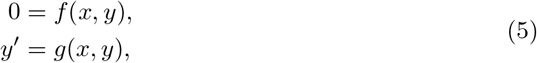

where ′ denotes the derivative with respect to the *slow* time *τ* = *ϵt*. Furthermore, for points in 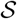 satisfying *∂_x_f*(*x, y*) ≠ 0 we know from the implicit function theorem that we can write a function *x* = *m*(*y*) from *f*(*x, y*) = 0, so we can express (5) as

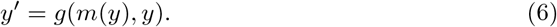

Therefore, once the trajectory has converged to the slow manifold, 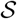, the *y* variable evolves following the dynamics in (6), while the *x* variable is given by *x* = *m*(*y*). So trajectories slowly move along 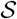 until reaching the fold points 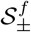. There, they become governed by the fast dynamics, and jump to the other stable branch. Indeed, as Fig. 1 shows, this is the mechanism underlying the generation of a stable periodic orbit Γ_*ϵ*_ showing relaxation oscillations, that is, the motion over Γ_*ϵ*_ consists of the alternation of long intervals of quasi-static behaviour (corresponding to the stable branches 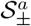 of 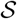) and almost instantaneous transitions between the branches [26].

**Fig 1.**
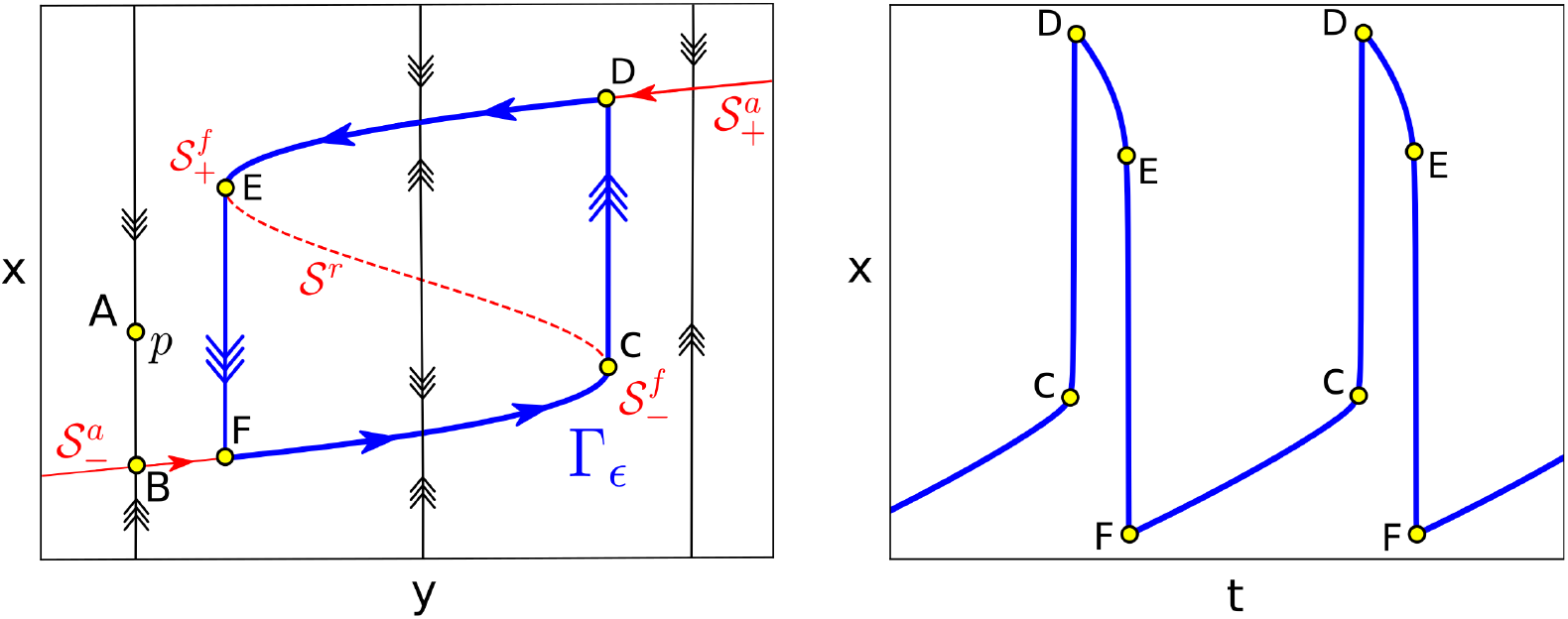
Phase space for relaxation oscillators. The slow manifold, 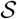, is a S-shaped curve having two stable branches 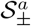 (solid red line) and one repelling 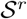 (dashed red line) (see Eq. (4)). Stable and unstable branches of 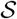 are separated by the fold points 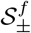. A given point, *p*, (see A) will quickly converge to the attracting branch of the slow manifold 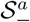 (see B). Then, it evolves along 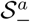 following (6) until reaching the fold point 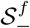 (see C) where it traverses fast to the other branch 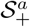 (see D). Then, following again the slow dynamics, the trajectory approaches 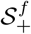 (see E) where it goes back to 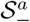 (see F). Therefore, the system (1) in the singular limit (*ϵ* → 0) admits a singular periodic orbit Γ_*ϵ*_ (in blue) generating relaxation oscillations.

### Phenomenological Epilepsy model

As we discussed in the introduction, the mechanism of relaxation oscillations (see Fig. 1) has been recently used in [17] to explain the apparent contradictory role of IEDs in epilepsy. In this work, the authors propose the following simple phenomenological epilepsy model, further referred to simply as the *phenomenor*:

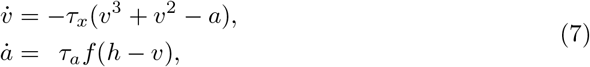

where *v* and *a* represent the firing rate and the excitability of a neuronal population, respectively. The dynamical changes in the excitability depend on the difference between *v* and *h* through the function *f*(*x*) = (tanh(*cx*) – *a*_0_), that is, an hyperbolic tangent whose slope is given by *c*. When *v* values are below *h*, the excitability increases, whereas when *v* values exceed *h* excitability decreases. Hence, *h* can be thought of as a threshold. For this study, *h* = *h_m_a* – *h_n_*. We keep fixed the particular set of parameters

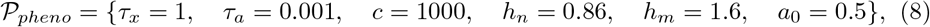

for which the system (7) displays a limit cycle denoted as Γ_*pheno*_ with a period of *T* ≈ 508.42; although the qualitative behaviour of the model stays the same for a wide range of parameters. Indeed, as *τ_a_* << *τ_x_* and the fast nullcline 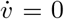 – which corresponds with the slow manifold 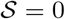 in (3) – describes a cubic curve, dynamics over Γ_*pheno*_ consists of a periodic switching between the states of low and high activity within relaxation oscillations.

The following Fig. 2 illustrates the mechanism proposed in [17] by which the phenomenor (7) reconciles the antagonistic role of IEDs. Consider the IEDs as a random train of pulses whose inter pulse interval distribution, *t_s_*, follows a normal distribution with mean value, *T_s_*, and standard deviation, *σ*: 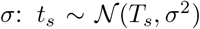. Whether or not a given perturbation causes an immediate transition to seizure depends on whether the perturbation manages to make the trajectory cross from the lower branch above the middle branch of the v-nullcline. If this happens, the trajectory rapidly converges to the upper branch, i.e. transitioning to the seizure regime. However, the response of the system dramatically changes depending on the amplitude, *A*, and mean inter pulse interval, *T_s_*, of IEDs (see panels A and B in Fig. 2).

The effect of a single pulse applied to the system, while on the lower branch, is either to keep the trajectory on the lower branch or to cause a transition to the upper branch. Therefore, the total effect of a train of pulses depends on the proportion of pulses causing transitions. Indeed, this dependence can be seen by plotting the change in the seizure rate Δ as a function of both the amplitude, *A*, and the mean inter-perturbation interval, *T_s_* (Panel C).

**Fig 2.**
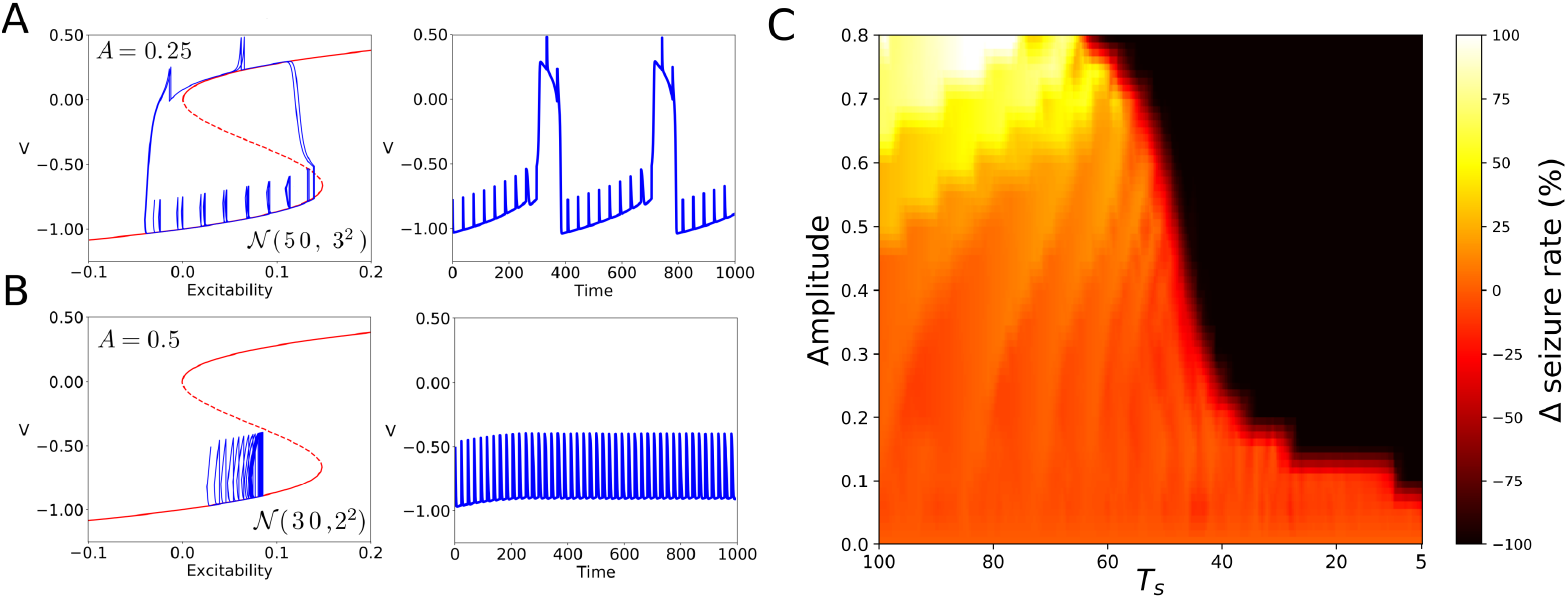
The antagonistic effect of IEDs on the transition to seizure. Panels A and B show, in red, the v-nullcline whose stable branches correspond to the stable low and high activity states of the system. The unstable part of the v-nullcline (dashed red line) separates the basin of attraction of both branches. Whether the pulses make the system cross the unstable part of the v-nullcline determines the opposite nature of IEDs. For a random train with amplitude *A* = 0.25 and 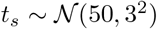 the system goes to seizure (panel A). By contrast, for a random train with amplitude *A* = 0.5 and 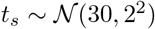 the system avoids the seizure state (panel B). By plotting the change in seizure rate Δ as a function of both the amplitude, *A*, and the mean inter-perturbation interval, *T_s_* (panel C). We can distinguish between pro-convulsive regimes (yellow and white areas) in which the transition is potentiated, and preventive regimes (red and black areas) in which the transition is delayed or completely suppressed.

### Phase Dynamics

Oscillations correspond to attracting limit cycles whose dynamics can be described by a single variable: the phase. As we now expose, the study of the dynamics on a limit cycle by means of the phase variable provides a more intuitive and simplified view of itssynchronization properties. Consider an autonomous system of ODEs

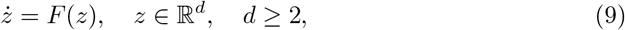

whose flow is denoted by *ϕ_t_*(*z*). Assume that *F* is an analytic vector field and that system (9) has a *T*-periodic hyperbolic attracting limit cycle, Γ. This *T*-periodic limit cycle, Γ, can be parametrized by the phase variable *θ* = *t*/*T* as

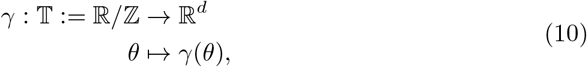

so that it has period 1, that is, *γ*(*θ*) = *γ*(*θ* + 1). While originally defined only on the limit cycle, the phase can be extended to the whole basin of attraction of Γ (which we will denote by 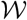). Indeed, as we consider attracting limit cycles, any point in 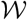 converges towards Γ as time tends to infinity. Therefore, we will say that two points *p* and 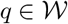 have the same asymptotic phase if

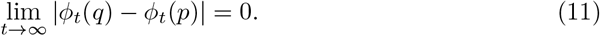

We further define the isochron 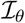 as the set of points having the same asymptotic phase *θ* [29], that is,

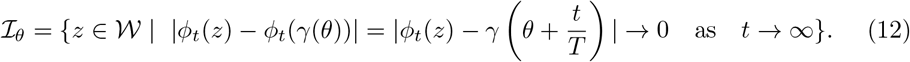

Let us now consider the effect of an instantaneous delta-like pulse over the *T*-periodic limit cycle Γ,

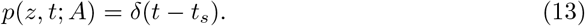

It is clear that the perturbation will just change the trajectory from one point *z* to another point 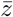. As we illustrate in Fig. 3, since the isochrons foliate the whole basin of attraction 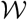 of Γ, we can say that the perturbation moved the trajectory from one isochron 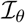 to another isochron 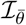, thus causing a phase shift 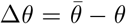. However, the phase shift will depend on the amplitude of the pulse and on the phase at which it was applied. This dependency is captured by the Phase Response Curves (PRCs). They are calculated by applying the same pulse to the limit cycle at different phases and registering how much the phase is advanced (or delayed). Let *z* = *γ*(*θ*) be a point on the limit cycle Γ. If we consider an instantaneous pulse as (13), it is clear that it will move *z* to 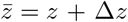. Thus, the PRC is defined as

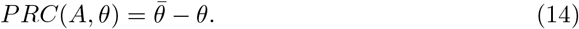

As Fig. 3 panel A shows, the isochrons 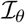 of Γ_*pheno*_ portrait the distribution of phases along the basin of attraction 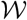. Whereas the isochrons for the upper branch of the cycle are almost vertical, the isochrons for the lower branch of the cycle show a more interesting geometry: they start vertical until crossing the *a*-nullcline, when they all bend. The shape of the PRCs as the amplitude, *A*, of the pulse increases is determined by this particular geometry of the isochrons. Since there is an almost constant distance of 0.1 between the lower branch of Γ_*pheno*_ and the slow nullcline 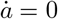, we can distinguish between two cases. For perturbations of *A* < 0.1, the perturbed trajectories only reach the part of isochrons consisting in almost vertical lines. Therefore, the corresponding phase shift Δ*θ* for perturbations on the upper and lower branches of Γ_*pheno*_ is almost negligible. Hence, the PRC for these phases will be close to zero. Indeed, only in the vicinity of the jumping points the PRCs will show larger values (see zoom window in Fig. 3C).

**Fig 3.**
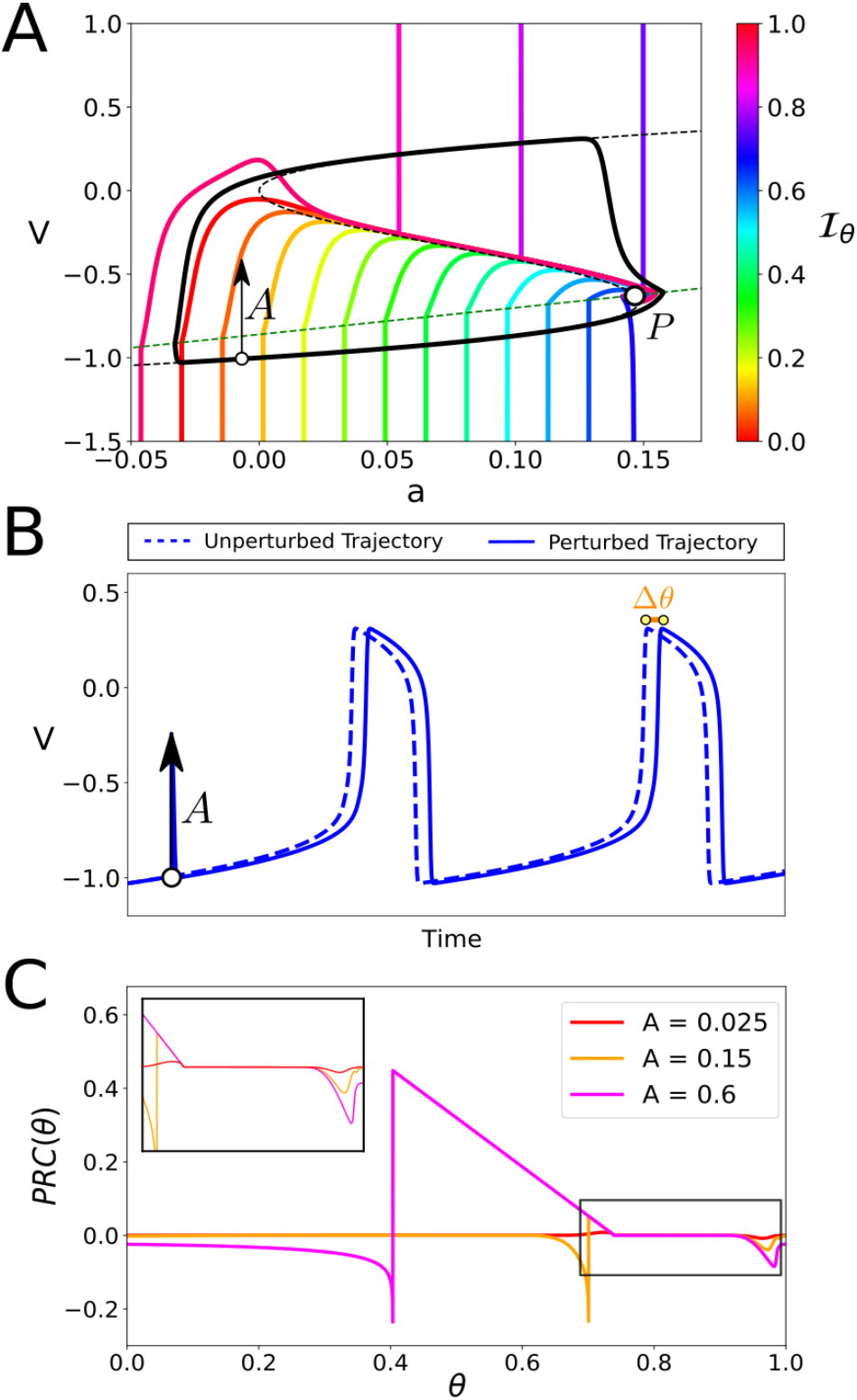
Isochrons and PRCs for the phenomenon. For the phenomenor (7) with the set of parameters 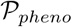 in (8) we show: (A) Limit cycle Γ_*pheno*_ and 16 equispaced isochrons 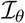 (left), the *v* and *a* nullclines (dashed black and green curves, respectively) and the fixed point, *P*, at their intersection. As panel (A) shows, since the isochrons foliate the whole phase space, a pulse amplitude, *A*, displaces the trajectory from an isochron 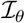 to another isochron 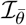 thus causing a phase shift 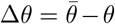 between the perturbed and unperturbed trajectory (see panel B). By applying the same pulses for all the points (phases) at the cycle and computing its respective phase shift, one computes the PRCs. The panel (C) shows the PRCs for the phenomenor for positive voltage pulses of different amplitudes.

By contrast, for perturbations of *A* > 0.1, the change on the geometry of isochrons for points on the lower branch remarkably changes the shape of PRCs. Perturbations on the lower branch will have a delaying effect unless they bring trajectories above the middle branch of the v-nullcline – which corresponds with 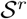 in (4) – so they advance phase. The delaying or advancing effect of a given pulse of amplitude, *A*, is delimited across a discontinuity for its corresponding PRC at the exact phase *θ*\* for which 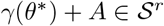.

The isochrons and PRCs computed for the phenomenor provide insight about how the combination of both the amplitude, *A*, and the mean inter pulse interval, *T_s_*, generate the different seizure propensity regimes in Fig. 2C. As isochrons in Fig. 3 show, positive voltage pulses of amplitude, *A*, at a point, *z* = *γ*(*θ*), on the lower branch cause a delay Δ*θ* < 0. However, for large enough mean inter-pulse intervals *T_s_*, although perturbations delay the system, they are not frequent enough to stop it from eventually transitioning to seizure (see Fig. 2 panels AB). Moreover, larger pulses are able to cause the trajectory to cross the v-nullcline earlier through the cycle (way before the fold point). Thus, the larger the amplitude of the pulse, the more common are these transitions.

By contrast, for small enough inter-pulse intervals, *T_s_*, the transition to seizure can be delayed or even stopped across the accumulation of the delays caused by each single pulse. Thus we can conclude that the mechanism underlying the description of the phenomenor of the role of IEDs, relies on the one hand on its cubic v-nullcline structure, allowing for relaxation oscillations and on the other hand on the prevalence of delays for positive perturbations at points on the lower branch not crossing the middle branch of the v-nullcline.

### Phase analysis of relaxation oscillators

As explained in the previous Section, the accurate description of the role of IEDs provided by the phenomenor is based on the prevalence of delays for perturbations in the ‘non-epileptic’ state, i.e. on the bottom branch of the cycle Γ_*pheno*_. Since this determining feature of the model - the prevalence of delays - is based on the bending in a particular direction of the isochrons, we aim to identify which elements in the model are key to cause this particular isochron geometry. As we show next, we perform this identification by taking advantage of the dynamical properties underlying any relaxation oscillator.

#### The 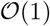 geometry of isochrons 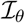

Next, we discuss some generalities shaping the isochrons of planar relaxation oscillators. To begin, it is worth recalling that if two points 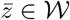 and *z* = *γ*(*θ*) belong to the same isochron, 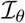, they have to meet at the same point of the cycle after a large enough time, *t* (see Eq. 12). For this reason, the determination of the shape of isochrons requires to study the converging dynamics towards Γ_*ϵ*_ which we recall that will consist of trajectories covering 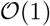 distances in the fast direction and 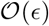 distances in the slow direction.

Since we aim to study the isochrons for relaxation oscillators, we can take advantage of the time-scale separation to be more precise concerning this convergence. Consider a point 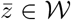. In a first approximation one can assume that the convergence of 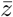 is achieved simply following the layer dynamics (2). If that was the case, since the layer dynamics consider the variable *y* as frozen, the isochrons will always be lines of *y* constant that we denote as 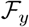. However, for correctly determining the shape of isochrons, we have to take into account that neither the convergence towards the limit cycle Γ_*ϵ*_ is instantaneous nor the dynamics on *y* during convergence are negligible. As a result, the isochrons are expected to be 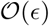 corrections of 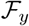. Indeed, it is worth to note that generalities determining the sign of those 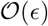 corrections will explain the prevalence of delaying (or advancing) effects of delta-like pulses in the fast direction.

Regarding the time needed for solutions to converge to the limit cycle, although the convergence towards a normally hyperbolic attracting limit cycle is ensured [30], for the case of slow-fast dynamics we can give even more details about this convergence by means of Tihonov’s theorem [31] (see also [32, 33]). Roughly speaking, Tihonov’s theorem states that after a time 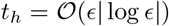, all orbits starting in a neighbourhood 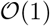 of the slow manifold 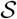 will have reached a neighbourhood of 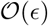 of 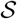.

Once we know the time, 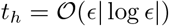, needed to converge, we can compute the motion of the converging point 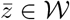 in the slow direction. The travelled distance in the *y* direction by 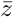 to approach a 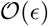 neighbourhood of Γ_*ϵ*_ is given by

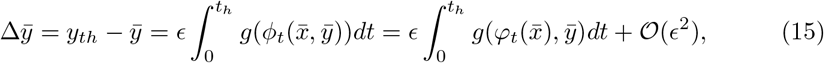

where 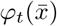 refers to the solution of the layer system (2). During the time, *t_h_*, needed to converge, the point *z* = *γ*(*θ*) on the limit cycle has travelled a distance Δ*y* given by

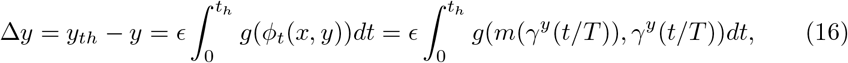

where in the second equality we utilize the fact that for points on the slow manifold we can use Eq. (6).

Now, let us illustrate how the difference between the distances travelled by the base point and the converging point, Δ*y* – Δ*ȳ*, will determine the sign of the 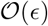 correction for isochron 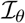 at the point 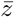. If we write the expression for Δ*y* – Δ*ȳ*:

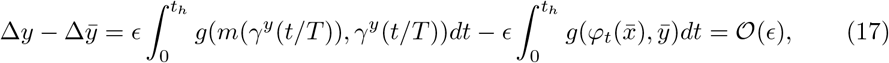

we can see that the difference Δ*y* – Δ*ȳ* is directly determined by the difference between the speeds in the slow direction for the base *z* and converging 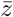 points during the time 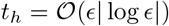 needed for approaching Γ_*ϵ*_. Basically, since both points *z* and 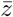 have to meet at the same point after the same time, the one travelling slower, needs to travel less distance. The difference, Δ*y* – Δ*ȳ*, corresponds exactly to the 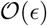 correction to 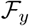 (see Fig. 4).

**Fig 4.**
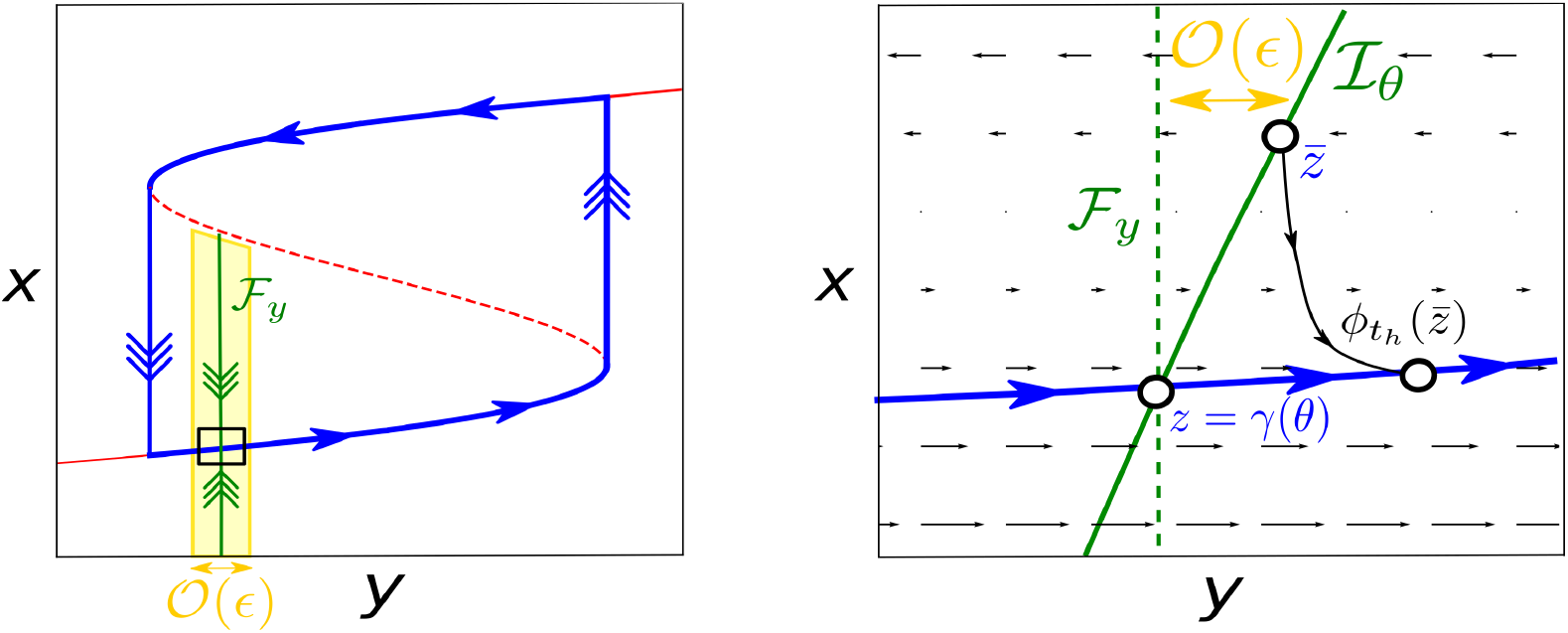
The slow vector field shapes the isochrons for relaxation oscillators. In the limit *ϵ* → 0 isochrons are lines of *y* constant denoted by 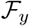. However, since *ϵ* ≠ 0 but small, the isochrons are 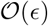 perturbations of 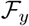. As we show in the right panel, the sign of the 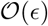 corrections depends on the difference of speeds between the converging point 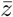 and the base point *z* during the convergence time *t_h_*. In this case, to approach Γ_*ϵ*_, 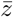 has to cross layers of *x* whose values are smaller than the ones surrounding Γ_*ϵ*_. For this reason 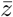 travels slower than *z*. Since 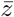 and *z* have to meet after a time *t_h_* at the same point on Γ_*ϵ*_, but 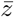 travels slower than *z*, then 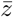 needs to travel a short distance. This determines the sign of the 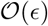 correction. Furthermore, if the slow vector field is monotonous along the fast direction, the farther the point 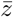, the slower (faster) it travels, so the slope of the isochrons will have the same sign for all the points 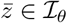 satisfying fast convergence, thus determining the effect of perturbations in the fast direction.

However, at the moment we have a local argument just justifying the shape of isochrons for a given point 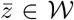. Nevertheless, we can globalize this argument by assuming some conditions for *g*(*x, y*). In particular, as we show now, if the slow vector field *g*(*x, y*) is monotonous in the fast direction, then the slope of the isochrons will have the same sign for all the points 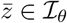 satisfying fast convergence.

The asymptotic phase defined in (11) allows to assign a phase to any point 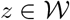, by defining the following function Θ(*z*)

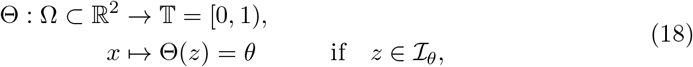

whose level curves indeed correspond to the isochrons. Let us assume we can invert Θ(*x, y*), so we can define the following function

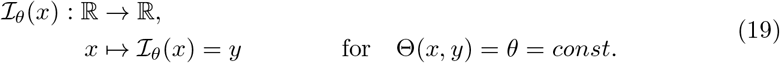

The slope of isochron 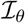, which we denote by 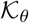, is then given by

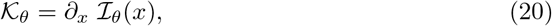

as we also have

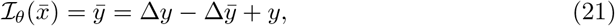

we can write the following expression for the slope 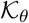

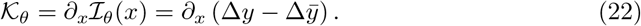

As the term Δ*y* – Δ*ȳ* can be written in integral form (see Eq. (17)), the slope 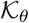, can be evaluated as the derivative of the difference of two sums (integrals)

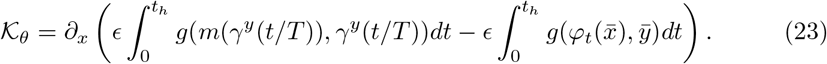

As we see, assuming that the vector field *g*(*x, y*) is strictly increasing (decreasing) function with *x* it is easy to discuss the sign of 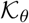. If the trajectory followed by the approaching point, satisfies 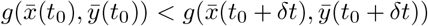 for 0 < *δt* ≤ *t_h_*, then, the second integral will be smaller than the first one. Since this difference will increase with 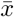, then 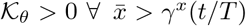. Furthermore, the larger the changes in *g*(*x, y*), the larger the slope. We remark that in the case 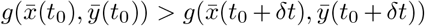, we can argue identically to obtain 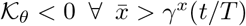.

In conclusion, we have illustrated the relationship between geometry of isochrons for relaxation oscillations and the slow vector field. First, we have shown how the tilt of the isochron 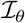 at a given point 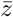 depends on the difference of speeds between 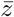 and the base point *z* during convergence. Furthermore, we showed that if the monotonicity of the vector field does not change, the tilt of the isochrons does not change sign as well.

We can illustrate these theoretical results by revisiting the isochrons for the phenomenor. As Fig. 5 shows, the parameters 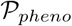 in (8) were chosen so that the tanh in (7) acts almost as a step function. As a result, the speed in the slow direction dramatically changes when crossing the slow nullcline. Since there are almost no differences between speeds for points below the slow nullcline, the isochrons are almost vertical. By contrast, this large difference of speeds once the slow nullcline is crossed, results in a remarkable bending of the isochrons for points on the lower branch of Γ_*pheno*_.

#### PRCs

Since the shape of PRCs is determined by the geometry of isochrons, next we discuss the extensions of our previous analysis of isochrons to PRCs. First, we can consider the limit *ϵ* → 0. In this case the isochrons would be vertical lines. Therefore, for points in the lower branch, unless the pulse brings trajectories above the mid-branch 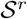 of the slow nullcline, its corresponding phase shift will be zero. For those points going to the other branch, the phase shift will be proportional to the skipped segment of the cycle, thus generating the characteristic shape of PRCs for relaxation oscillators [34] (see black curve in Fig. 6 right). However, our knowledge of the geometry of isochrons can extend this result. Without loss of generality we discuss the case 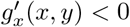. In this case, we know that perturbations acting over points on the lower branch not crossing 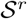 will delay the system (see Fig. 4). As a consequence, the PRC will have negative values for all the phases *θ* in the lower branch such that *θ* < *θ*\* where 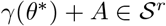. Although the particular shape of the delaying segment of the PRC will depend on the particular slow vector field chosen, in general, we expect the crossing of the slow and fast nucllines to generate a single unstable fixed point (denoted by *P*) inside Γ_*ϵ*_. It is worth to mention that since isochrons will approach *P* through 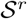 [35], we expect the bending of a particular isochron to increase as it approaches 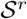. As a consequence, we expect the maximal delay values of a PRC to concentrate near the jumping phase, *θ*\*. Finally, if we consider perturbations over points in the upper branch, arguing similarly as in Fig. 4, we can conclude that the effect of pulses of positive amplitude is to advance trajectories.

**Fig 5.**
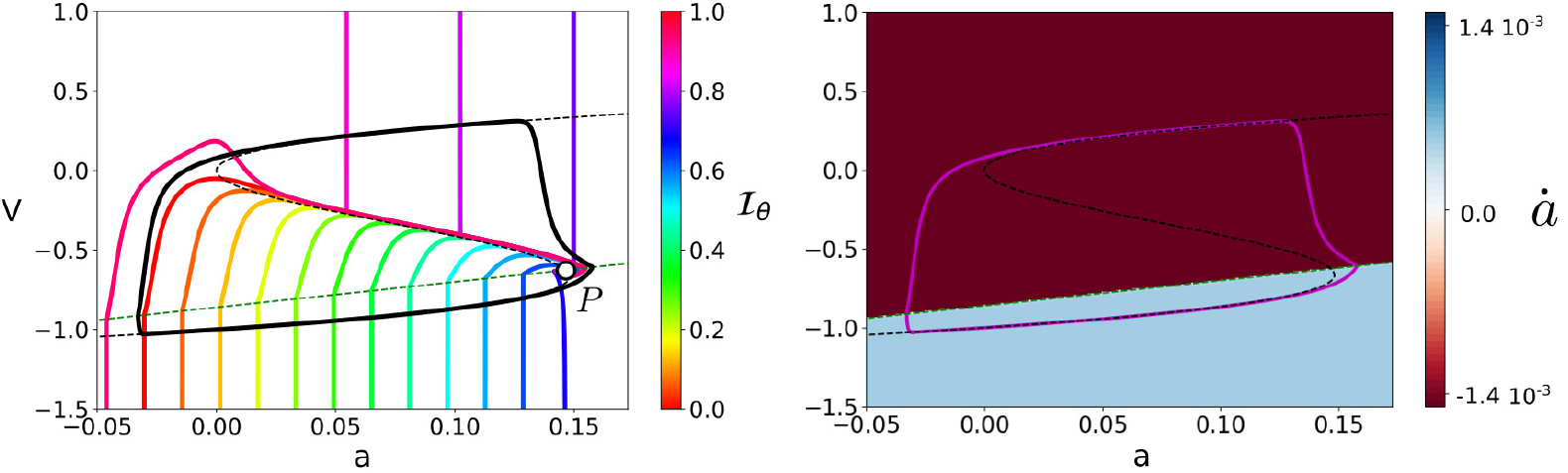
Relationship between the curvature of isochrons and the values for the slow vector field. For the phenomenological epilepsy model (7) with the set of parameters 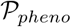 in (8) the figure shows: (A) Limit cycle Γ_*pheno*_ and its isochrons 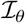 (left). (B) Values of the slow vector field (corresponding to 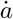 in (7)) for points 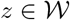.

**Fig 6.**
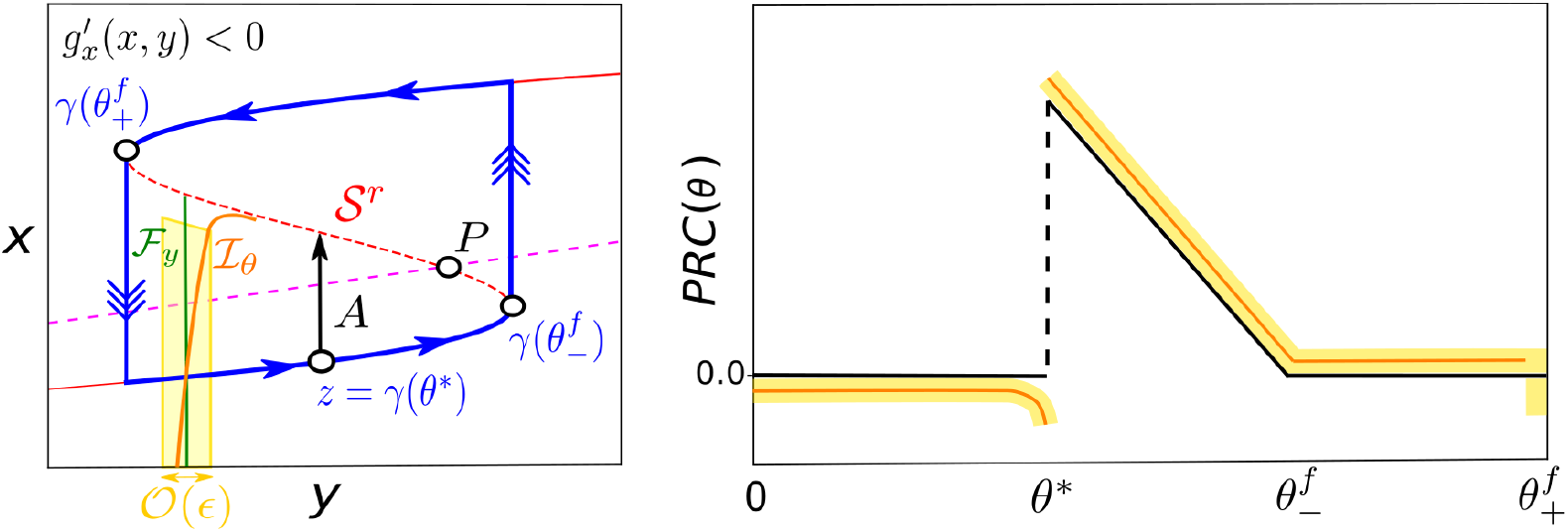
PRC of pulses *A* > 0 for relaxation oscillators. Next we sketch the PRCs for pulses of amplitude *A* > 0 for the case 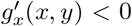. For phases *θ* < *θ*\*, where *θ*\* satisfies 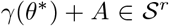, due to the slope of isochrons the effect of the pulses will be to delay trajectories. Since isochrons approach the unstable point *P* through 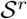, the near the phase *θ* to *θ*\*, the larger the bending of the isochrons and thus the larger the corresponding delay value. For phases 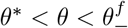, there is an advancement proportional to the fraction of cycle skipped. This prevalence of advancements is also seen for points in the upper branch. For phases in a neighbourhood of the fold point 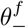, we expect a transition between advancement and delays not drawn because our analysis is only valid for normally hyperbolic points.

#### Phase locking

So far we have theoretically identified the factors shaping the isochrons for relaxation oscillators. Furthermore, we have discussed how the particular geometry of the isochrons for relaxation oscillators is reflected in the corresponding PRCs. Next, we aim to continue extending our theoretical approach to determine generalities underlying the mechanism by which external perturbations suppress the original oscillatory dynamics. We recall that, in the particular case of epilepsy, we are studying the suppression of the original oscillation through the accumulation of delays which causes the system to remain in the lower activity state and thus to prevent the transition to seizure.

A delta-like pulse of amplitude, *A*, reaching the cycle at a phase, *θ*, will map it to a new phase *f_A_*(*θ*) = *θ_new_*, where the map *f_A_*(*θ*) writes as

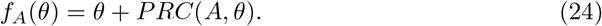

If the perturbation was a train of periodic pulses with an inter stimulus interval given by *T_s_*, we can describe the phase dynamics of the system by

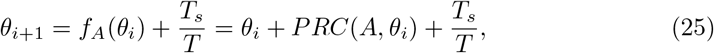

where *θ*_0_ = *θ*. The fixed points of the above map (25), which are given by

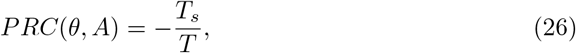

correspond to the phase locking states of the system.

As Eq. (26) shows, the PRC determines the asymptotic state of the perturbed dynamics. However, not all the phase locking states predicted by Eq. (26) correspond to the particular locking mechanism we are looking for. For example, if we consider very small delay values Δ*θ* → 0^−^, it is clear that values of *T_s_* ≈ *T* will correspond to a phase locking state of the system which does not prevent the transition to seizure. The particular mechanism we are looking for is depicted in Fig. 7. Consider a pulse displacing a point *z* = *γ*(*θ*) to 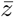. If we denote by *t_h_* the time that 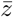 needs to approach Γ_*ϵ*_, we need 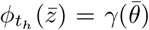 with 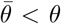. That is, we need the perturbed trajectory to reach the cycle at a previous phase. Assuming fast convergence, we can write

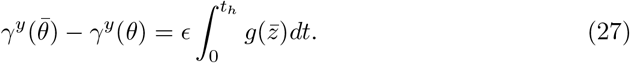

Since we need 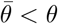 as a necessary condition for phase locking, then, if we assume without loss of generality that the motion over Γ_*ϵ*_ is counter-clockwise, the above integral has to be negative. For that to happen, the perturbation has to necessarily send trajectories above the slow nullcline. Indeed, if we denote the by *t*\* the time needed to cross the slow nullcline, then, the particular class of locking we are interested in has to satisfy

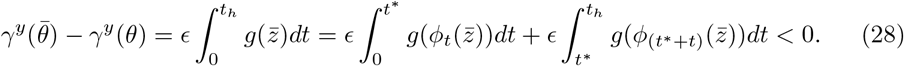

Since the first integral is negative and the second is positive, above Eq. (28) shows that the appearance of phase locking requires the perturbed trajectories to be sent to a point such that the distance travelled during convergence in the negative direction overcomes the distance travelled in the positive direction, so the total displacement is negative. Then there is a time *T_s_* = −*T*Δ*θ* > 0 (with 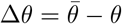) for which the next pulse will kick the system at the same initial point *z* = *γ*(*θ*) (see Fig. 7). The repetition of this process keeps the trajectory on the lower branch, and prevents the seizure emergence by suppressing the original oscillatory dynamics. Importantly, we highlight the strong influence of the slow vector field on the appearance of this locking mechanism. Indeed, the smaller the distance between the slow nullcline and the lower branch, the smaller the amplitude of perturbations needed for locking the system.

**Fig 7.**
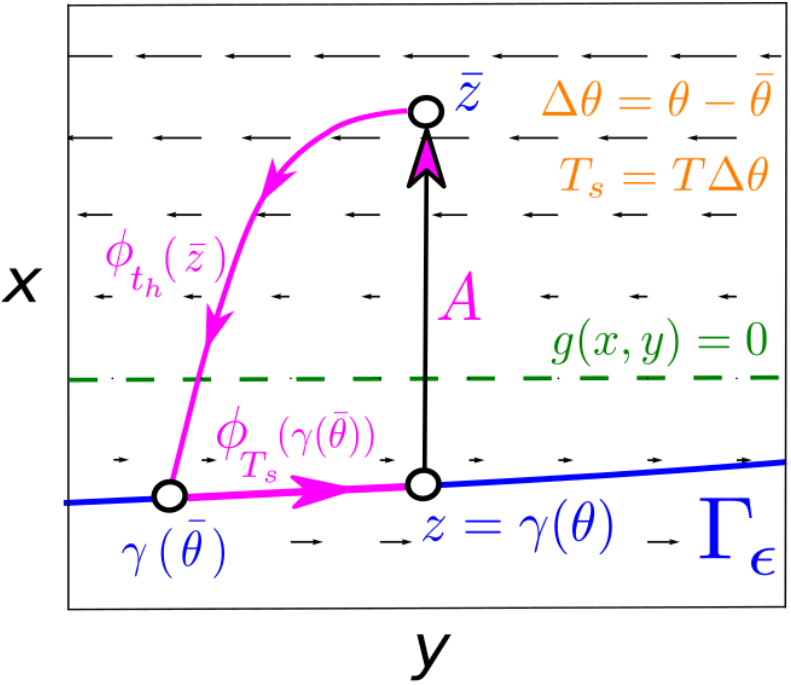
Mechanism preventing the emergence of seizures. To suppress the original oscillation and keep the system in the lower branch of Γ_*ϵ*_ the amplitude *A* of the pulse has to be large enough so besides causing a delay Δ*θ*, it displaces trajectories above enough the slow-nullcline so the distance travelled in the negative direction overcomes the distance travelled in the positive direction, thus causing a negative net displacement. The locking appears by repeating this mechanism after *T_s_* = *T*Δ*θ* intervals so the new pulse always hits the system at the same initial point.

We can check the validity of this result by revisiting the results for the phenomenor. Fig. 8A shows the relative seizure rate increase Δ due to a train of random perturbations for a *T_s_* periodic train of pulses. We can see how the locking preventing the transition to seizure starts for values *Ā* ≈ 0.1, which is the approximate distance between the lower branch and the slow-nullcline. Furthermore, for a fixed amplitude *A* > *Ā*, if we consider the maximum delay value (denoted by Δ*θ*\*) of the corresponding PRC and compute the inter pulse interval value given by 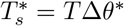, it is clear that for inter-pulse intervals 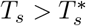, the system is likely to jump because the delays are not large enough to stop the system. Therefore, we expect the pair 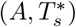 to delimit the locking regime. By computing the PRCs for all the amplitude values satisfying *A* > *Ā*, we can calculate the corresponding 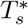 values and thus generate a curve in the (*A, T_s_*) space - which indeed corresponds with the bifurcation curve of the map (25) – showing a nice agreement with the boundaries of the locking area (see purple line in Fig. 8A).

Results for the random case in Fig. 8B can be interpreted by means of the periodic case. The random dynamics can be computed as well by using a similar map to (25) but substituting *T_s_* for *t_s_* values in the distribution 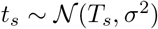. In this case, the system does not ‘lock’ in the same way the deterministic system does, that is trough fixed points in Eq. (26). However, one might try to interpret random dynamics by means of the periodic case. By computing the maximum delay value Δ*θ*\* of the PRC, we calculate the characteristic value of 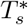, corresponding to the phase *θ*\* such that perturbations 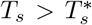 will jump. Therefore, the robustness of the deterministic locking states to noise, will be determined by whether 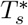 is or not into the width *σ* of the inter pulse distribution.

### Epileptor model

One of the most known and widely accepted models in epilepsy is the epileptor model [13]. This model consists of 5 differential equations (4 fast and 1 slow) so it can display a wide range of dynamical regimes explaining many different pathways to seizure [36]. In order to show the generality of the results derived from our theoretical approach and to demonstrate their consequences in models of epilepsy, we will study the following 2D reduction of the epileptor model [37]:

**Fig 8.**
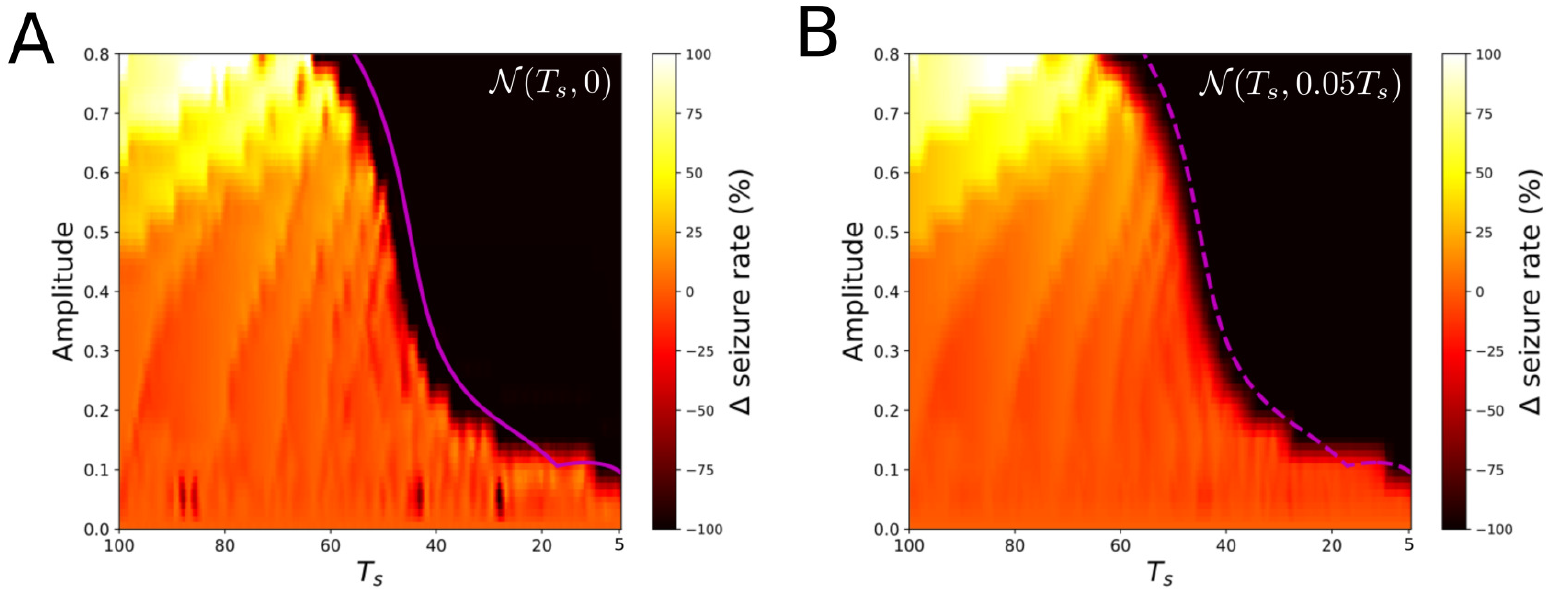
Response of perturbations for the phenomenor. We plot the change in the seizure rate Δ for a random train of pulses following a Gaussian distribution of mean time *T_s_* and standard deviation *σ* denoted as 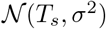. Panels (A) and (B) correspond to the deterministic periodic case 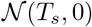 and to the random case 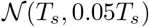, respectively. For panel (A) we plot a purple solid line corresponding to the bifurcation of the phase map (25). We plot the same curve as a dashed purple curve in panel (B) illustrating the resilience of the deterministic phase-locked states to noise.

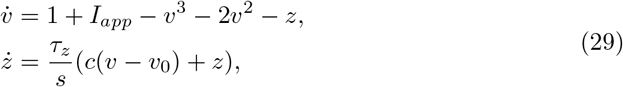

where *v* and *z* represent the firing rate and the permittivity of a neuronal population, respectively. For this model we will work with the sets of parameters 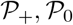 and 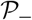 in Table 1.

**Table 1.** Different parameters for the reduced 2D Epileptor model in (29). For the set of parameters 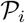, the system will display a limit cycle Γ_*i*_ of period *T_i_*.

| | $\tau_z$ | $x_0$ | $I_{app}$ | $c$ | $s$ | Lim. Cycle | Period |
| --- | --- | --- | --- | --- | --- | --- | --- |
| $\mathcal{P}_+$ | 1/2857 | -2 | 3.1 | -4 | -1 | $\Gamma_+$ | $T_+ \approx 2181.6$ |
| $\mathcal{P}_0$ | 1/2857 | -1.5 | 3.1 | -16 | -1 | $\Gamma_0$ | $T_0 \approx 695.7$ |
| $\mathcal{P}_-$ | 1/2857 | -0.1 | 3.1 | 2.4 | 1 | $\Gamma_-$ | $T_- \approx 7333.3$ |

Identically as the phenomenor, since the time constant for the *z* variable is small *τ_z_* << 1, and 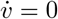 describes a cubic curve, the three sets of parameters 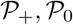 and 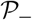 lead to relaxation oscillators denoted as Γ_+_, Γ_0_ and Γ_−_ respectively. The three different sets of parameters 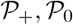 and 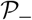 were chosen to illustrate the influence of the slow vector field on the response of perturbations of the system. Indeed, we denoted the parameters as 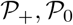 and 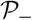 because they set the nullcline to have positive, horizontal and negative slope, respectively. Fig. 9 shows the isochrons and PRCs for the three sets of parameters 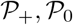 and 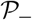. Since the slow vector field of the reduced epileptor is monotonic in *x*, the slope of the isochrons does not change sign for any of the considered cases, and again, it causes a prevalence of delays for perturbations of positive amplitude over points on the lower branch which is captured by the PRCs (see Fig. 9). We remark the similarity between the computed PRCs in Fig. 9 and the ones sketched in Fig. 6.

**Fig 9.**
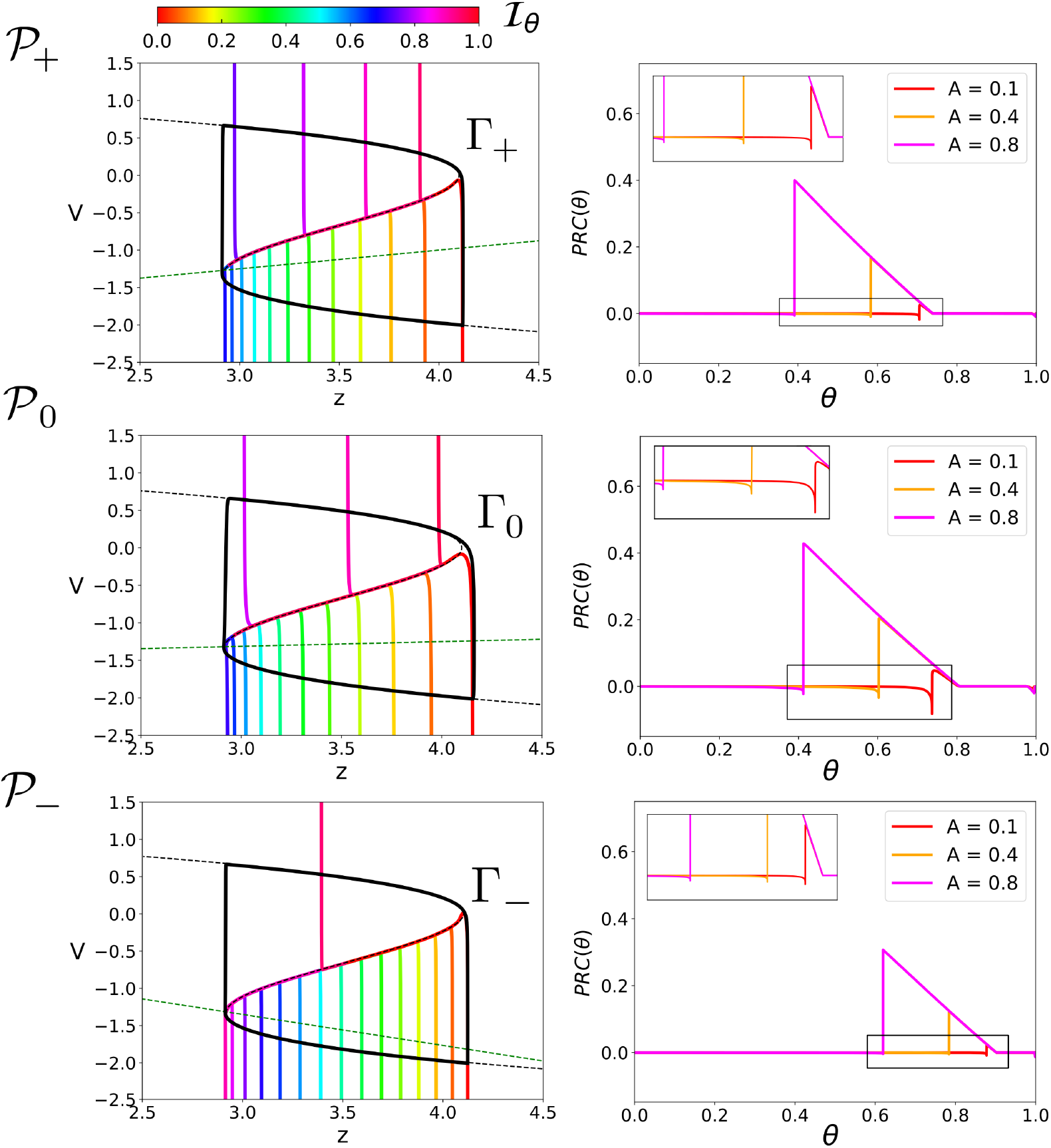
Isochrons and PRCs for the reduced epileptor. For the sets of parameters 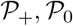 and 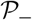 in Table 1 we show: Limit cycle Γ_+_, Γ_0_ and Γ_−_ and its isochrons 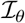 (left). The phase response curves in the *v* direction for different values of *A* (right). For the three cases we plot 16 equispaced isochrons. Consistently with our previous analysis, since the monotonicity of the slow vector field does not change, the slope of isochrons does not change sign.

### Response to perturbations

Next, we show how while the unperturbed behaviour of the cycles Γ_+_, Γ_0_ and Γ_−_ remains qualitatively identical, that is, they show relaxation oscillations, their response to the same train of pulses will be completely different. As we will argue, these remarkable differences can be explained by the different sets of parameters 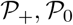 and 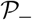 causing different slow vector fields for each cycle. Identically as in the phenomenor case, we consider a random train of pulses whose inter pulse interval follows a normal distribution of mean *T_s_* and standard deviation *σ*, denoted by 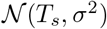 and compute the change of the seizure rate Δ for a train with 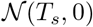 and 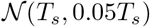.

The simulation results are summarized in Fig. 10. Consistently with the theoretical results, we can see a direct correspondence between the mean distance between the lower branch and the slow nullcline and the appearance of areas suppressing the oscillation. For this reason, Γ_−_ locks for smaller amplitude values than for Γ_0_ and Γ_+_. Furthermore, although the bending of the isochrons is small and so are the corresponding delays Δ*θ*, because of its large *T* value (see Table 1), the range of *T_s_* = −*T*Δ*θ* values for which Γ_−_ shows locking is even larger than for Γ_0_ and Γ_+_. We also remark the good agreement between the bifurcation curves of map (25) and the areas suppressing the transition to seizure.

Regarding the interpretation of the random perturbation train scenario, we can interpret results approximately by means of the results for the periodic perturbation scenario. Similarly as we argued in the phenomenor case (see Fig. 8), the robustness of a given locking state to noise will depend on whether the critical value of 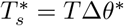, (where Δ*θ*\* corresponds with the maximal delay value of the PRC) is or not within the width *σ* of the distribution 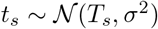. The higher the probability of occurrence of 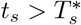 values, the likely is the system to switch to the upper branch. The differences in the resilience of the deterministic locking areas for Γ_+_, Γ_0_ and Γ_−_ in Fig. 10, can be explained by the different values of the period for the 3 cycles (see Table 1). Despite the PRCs for the three cycles show a similar range of values for the delays Δ*θ*, the differences come when these delays are transformed in inter impulse intervals through *T_s_* = *T*Δ*θ*. The shorter the period *T*, the smaller the critical 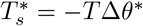 value. Since in the three cases the *t_s_* distributions have the same width, the smaller the critical 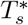 value, the higher the probability of occurrence of 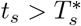 values. As a consequence, the resilience of locking states for Γ_+_ and Γ_0_ is weaker than for Γ_−_ in which the distribution *T_s_* = −*T*Δ*θ* is larger because of its larger period.

### Comparison between the phenomenor and the reduced epileptor

Although both the phenomenor in Eq. (7) and the reduced epileptor in Eq. (29) model seizure dynamics through relaxation oscillations, it is worth to mention the different role of the slow variable in the models. In the phenomenor the variable *a* describes the excitability of the tissue (the higher excitability, the more likely the spontaneous seizure initiation), whereas in the (both original and reduced) epileptor the *z* variable (dubbed as *permittivity*) has the opposite polarity: for its *low* values, the system switches to seizure as its only stable state. As a consequence, although the dynamical mechanism of the two models generate is virtually identical, the monotonicity of the slow vector field and the rotation direction over the cycle is flipped (see Fig. 11). However, in both models, the motion and the tilt of the isochrons are related in such a way that the prevalent effect of positive voltage perturbations over the lower branch of the cycle is to slow-down the oscillations, or in particular to delay the seizures.

From a mathematical perspective, the main differences between both models rely on their different time constant *τ* values and the specific slow vector field functions *g*(*x, y*). Because of the correspondence between *τ* << 1 and *ϵ*, we expect the isochrons to be bounded in domains 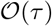 (see Fig. 4). However, from our analysis it also follows that the bending of the isochrons, although being contained in 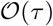 domains, will be also determined by dependence of *g*(*x, y*) on the fast variable *x* between the perturbed and the base trajectories (see Eq. (23)). To illustrate these role of *τ* and 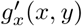, let us compare Γ_*pheno*_ with Γ_−_. In both cases, the slow nullcline was near the lower branch, so we have a (qualitatively) similar geometry for both phase spaces. For this reason the response to perturbations was qualitatively similar in both cases (compare Fig. 8 and Fig. 10C). However, the larger range of *T_s_* values for which perturbations over Γ_*pheno*_ avoid seizure can be explained by both the larger *τ* and the strong change in the monotonicity of *g*(*x, y*) for the PE. The combination of both effects causes a larger bending of the isochrons and thus larger delays. Therefore, for (qualitatively) similar geometries, the differences in both the time constants values and in the strength of variations in the fast component of the slow-vector field have a substantial effect on the amplitude of the phase response of the system to inputs.

**Fig 10.**
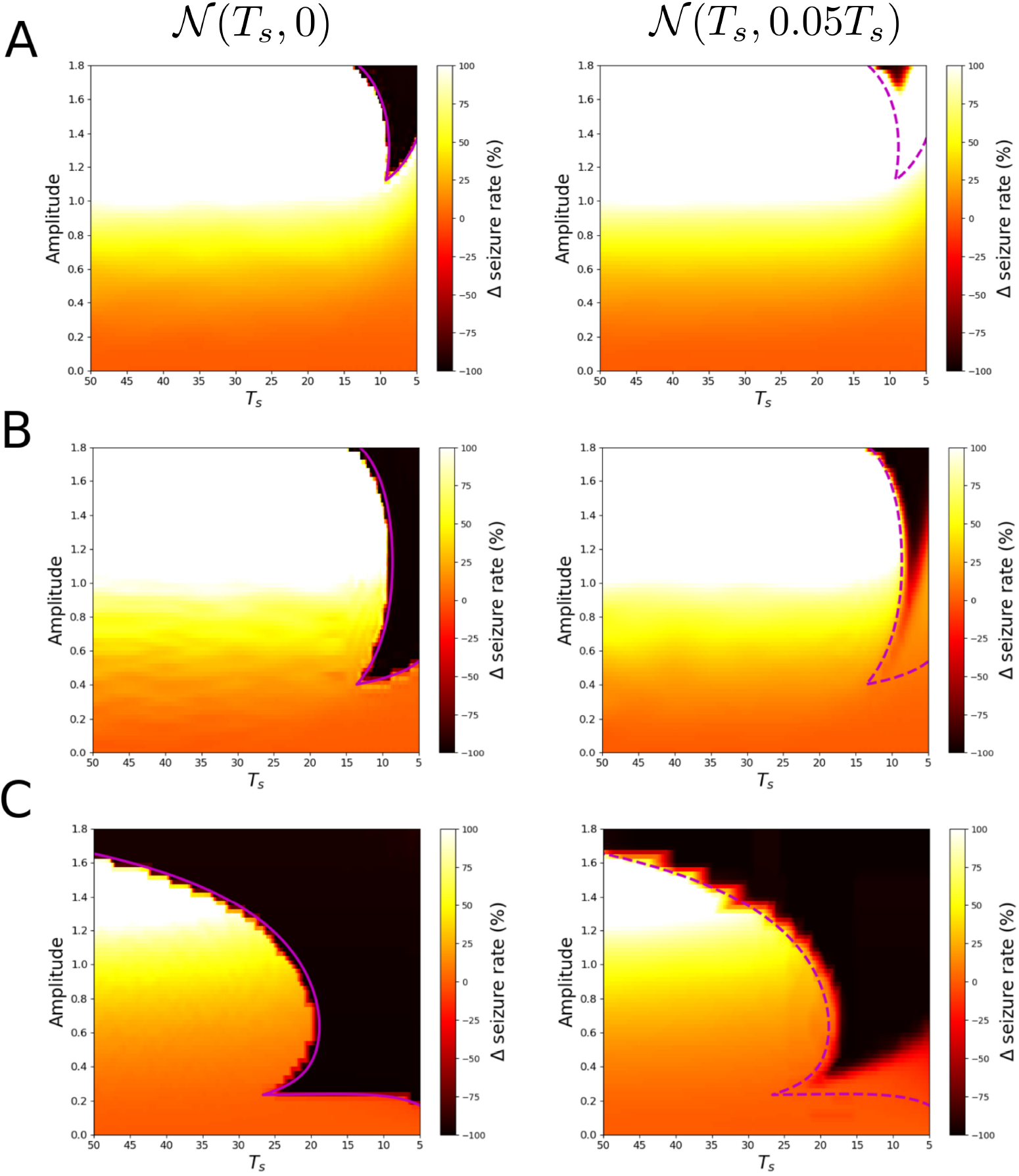
Response of perturbations for the reduced epileptor (29). We show the change in the seizure rate Δ for a random train of pulses whose mean inter impulse interval follows a normal distribution 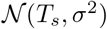 with a mean time *T_s_* and standard deviation *σ*. Panels A, B, C correspond to the sets of parameters 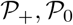 and 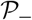 in Table 1. Left Figures correspond to the periodic case 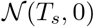 and right figures to the random case 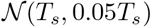. Consistently with our theoretical analysis there is a direct correspondence between the mean distance between the lower branch and the slow nullcline and the minimal pulse amplitude A for which perturbations may lead to lock the system. Purple solid lines, bounding locking regimes, correspond to the bifurcations of the map (25). By drawing the same curve for the random case, we illustrate the resilience of locking states to noise.

**Fig 11.**
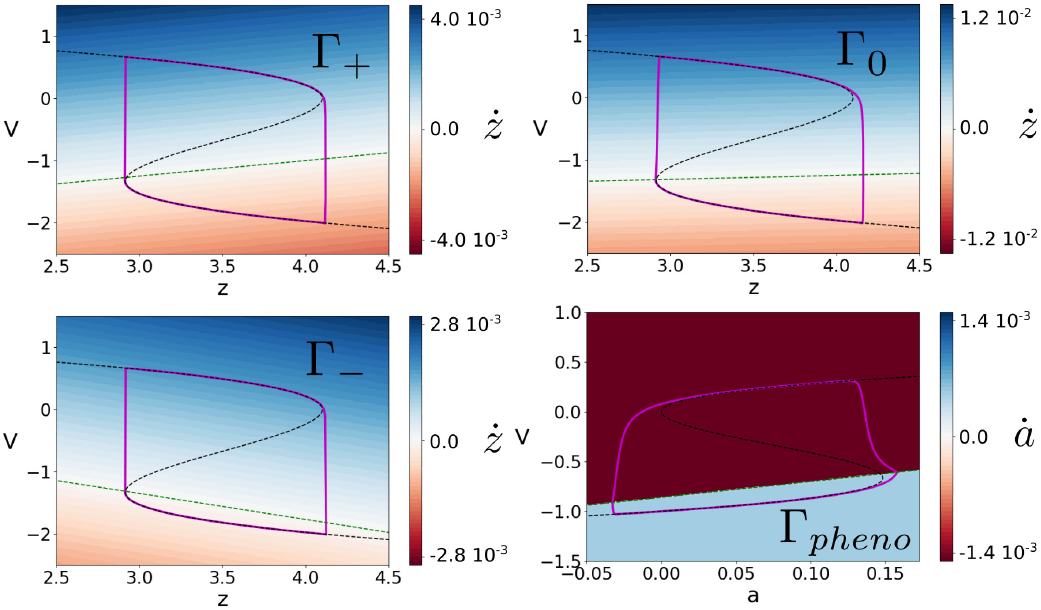
Slow vector field for the reduced epileptor and the phenomenor. Each cycle is depicted in purple, the v-nullcline in black and the slow nullcline in green. Notice that the direction of the slow variable in both models is flipped, and thus is also the motion over the cycles and the sign of 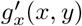.

## Discussion

In this paper we applied a phase approach to analyse planar relaxation oscillators, motivated by models of epileptic dynamics. Indeed, the study of neural oscillators by means of the phase reduction has been extensively utilized in neuroscience from the level of single neurons to the network scale [23, 38–40]. In this work, the computation of isochrons and PRCs of the phenomenological seizure dynamics model introduced in [17] fully clarified the mechanism integrating the antagonistic potential effects of IEDs. Furthermore, the theoretical analysis of the phase response of a generic planar relaxation oscillator manifested the crucial role of the slow vector field on the geometry of their isochrons. Due to the direct link between isochrons and PRCs, we have been able to study the relationship between the slow vector field and the different response behaviour a planar oscillator can display depending on the amplitude and frequency of perturbations. For the cases considered, whereas the distance between the slow nullcline and the bottom branch of the cycle indicated the minimum value of amplitude values for suppression of the original oscillation, the minimum value of PRCs (that is, the maximum delay) was related to the maximum interpulse intervals for which this locking mechanism holds. Furthermore, besides confirming our results, the study of variants of the reduced epileptor model [37] showed how vastly different responses to perturbations can be exhibited by models differing only in the slow-nullcline position, but possessing almost identical unperturbed behaviour, i.e. equivalent limit cycle oscillations, thus demonstrating the key role of the slow vector field in the response of perturbations for planar relaxation oscillators.

We acknowledge that due to the motivation by models of epilepsy, we showcased the theory only on a small set of example dynamical systems previously used for modelling the cyclical transition between an ictal and interictal state, which showed quite similar dynamics, including having one linear and one cubic nullcline, and a monotonous slow component of the flow field. A quick glance at other slow-fast relaxation oscillator models however suggests, that these properties are far from uncommon in many other models. Moreover, careful consideration of the theoretical arguments however shows, that the specific linear or cubic shape is indeed not crucial for the general observations to hold. Also, careful consideration of the theoretical arguments shows, that the monotonicity of slow vector field is firstly quite natural (the function needs to change from positive to negative values between the two stable branches of the stable manifold; the Occam’s razor suggests that it will likely do so monotonically); and moreover not necessarily needed - if the change is not monotonic, the dependence of the PRC on the size of the perturbation just becomes more complicated, however the (sign of) the PRC is still given by the integral of the slow component along the recovery trajectory.

Another apparent limitation is that we focused on the effect of positive pulses acting on the bottom branch of the cycle. However, the approach straightforwardly extends to planar oscillators having more complex slow vector fields and to pulses of different sign applied either to the lower or higher branch. Indeed, we suggest that for a given slow vector field the applied geometrical approach is instrumental in providing an intuitive insight concerning the isochrons and therefore the PRCs. In that sense, our analysis extends previous results on PRCs and isochrons of planar relaxation oscillators beyond the weak and singular limit [34, 41]. Theoretically more interesting, while also more demanding, is the generalization to higher dimensional oscillators, providing richer geometrical structure of the flow, perturbations and trajectories. However, previous simulation-based results on the full Epileptor model [17] suggest that the potential dual effect of perturbations on oscillatory behaviour is preserved even in higher dimensions, although richer behaviour might show for other models or perturbation scenarios.

Regarding epilepsy, our results indicate the key influence of the slow vector field on the propensity for seizure emergence. We acknowledge our analysis relied on reduced planar models. However, we plan to make advantage of recent methodologies computing isochrons of high dimensional systems [42] to extend our approach to different high dimensional models as [13, 15, 16, 43, 44]. In general, the high dimensionality of these models permits to describe more accurately seizures initiation and termination [12, 45]. We believe the continuation of this line of research may provide an alternative vision to the questions these models approach. Furthermore, because of the usage of the phase variable and the determination of PRCs, we think this approach can also help to determine more accurately coupling functions for studies approaching epilepsy from the coupling of different oscillatory units [46].

Importantly, the quest for deeper and intuitive understanding of the effect of perturbation on epileptic network dynamics is not a just an intriguing mathematical exercise, but an indispensable part of an important while difficult journey to understand the mechanisms of seizure initiation, and the possible ways to preclude this initiation by therapeutic stimulation interventions [47]. Of course, while the general conceptual insights are on their own relevant for general understanding the possible dynamical phenomena in response to perturbations, the observed role of the slow component of the field and in particular the nullcline suggests that any computational models of epilepsy dynamics should also attempt to reasonably approximate these aspects (and not only the unperturbed behaviour), if aspiring for providing relevant predictions concerning treatment protocols or just outcomes of endogenous perturbations and inter-regional interactions. This opens also the question of how to practically estimate these properties from experimental data, be it through stimulation protocols or purely observation data; this seems to be a natural avenue for obtaining more realistic models of epileptic dynamics.

In conclusion, we have outlined and carried out phase response analysis of planar relaxation oscillator models of epileptic dynamics that opens not only a path in epilepsy research with many interesting analytical, computational, experimental and potentially clinical implications, but also provides a framework applicable to gain insight in the plethora of other computational biology problems in which slow-fast relaxation oscillator models are pertinent.

## Materials and methods

This section contains some technical details concerning the numerical implementation of computations used to provide the presented results. Integration of ordinary differential equations was done using a 8th-order Runge-Kutta Fehlberg method (rk78) with a tolerance of 10^−14^.

### Computation of Isochrons

To compute isochrons of slow-fast systems, we start by computing the parameterization *γ*(*θ*) of Γ (see Eq. (10)). To do so, we construct a Poincaré section and use a Newton method to find a fixed point of the corresponding Poincaré map. By doing this, we obtain a point *x*_0_ ∈ Γ and the period *T*. Then we integrate the system (9) with initial condition *x*(0) = *x*_0_ for a time *T* to obtain *x*(*θT*) =: *γ*(*θ*) for *θ* ∈ [0,1).

Next, we need to compute the linearization *N*(*θ*) of the isochrons around Γ. To that aim typically one solves a variational-like equation [48]. However, in slow-fast systems the cycle is very strongly attracting (indeed, its Floquet exponent is 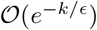 where *k* > 0) [49]. For this reason, obtaining *N*(*θ*) via numerical integration requires to deal with very small numbers, so one needs high precision algorithms and large number of decimals.

As an alternative to numerical integration we took advantage of the fact that ∇Θ(*x*) is perpendicular to the level curves of Θ(*x*), which indeed correspond to the isochrons. Therefore, we can use the infinitesimal PRC (iPRC), that is ∇Θ(*γ*(*θ*)), to compute *N*(*θ*) through the following equation [48]:

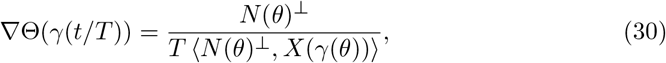

where *v*^⊥^ refers to a perpendicular vector to *v* and < ·, · > to the usual dot product. Instead of computing the iPRC ∇Θ(*γ*(*t*/*T*)) by integrating the adjoint equations (which also display numerical instabilities) we compute it by means of the procedure described in next subsection.

Finally, we globalise the isochrons via the backward integration of *N*(*θ*) (we refer the reader to [48] for more details about the globalisation procedure).

### Computation of PRCs

The PRCs in this paper were computed using a continuation method. The computation of PRCs by direct integration of the perturbed trajectories, usually measures the phase shift over the maxima of a certain variable. That is, they require a relaxation time *T_rel_* large enough so the perturbed trajectories reach the maximal values over the cycle. By contrast, as we now show, continuation methods just require the perturbed trajectories to reach a point on the cycle. Therefore, one needs to integrate a shorter time *T_rel_*. Specifically in slow-fast systems in which the periods of the system are large, the usage of continuation methods saves a lot of computational effort. To compute PRCs, we have used the continuation method introduced in [50], which we now briefly review for the sake of completeness.

A pulse acting on a point *z* = *γ*(*θ*) ∈ Γ will displace the trajectory to 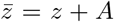. Then, after a time *T_rel_* large enough, the trajectory will be again on the limit cycle but with another phase 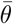. Mathematically

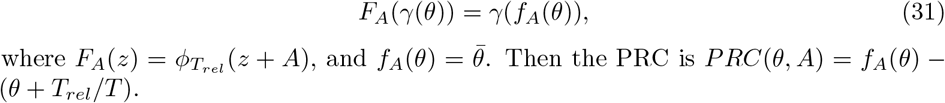

where *F_A_*(*z*) = *ϕT_rel_*(*z* + *A*), and 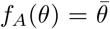. Then the PRC is *PRC*(*θ, A*) = *f_A_*(*θ*) – (*θ* + *T_rel_*/*T*).

The idea of the method is to obtain *f_A_*(*θ*) by solving Eq. (31). To that aim, one can use the following algorithm which computes the PRC for a perturbation of amplitude *A* by means of a Newton method. The computation of PRCs via continuation is achieved using the computed PRC as an initial seed for computing the PRC for a new amplitude *A*′ = *A* + Δ_*A*_. Given the parameterization of the limit cycle *γ*(*θ*), and *f_A_*(*θ*) an approximate solution of equation (31), we perform the following operations:

1. Compute *E*(*θ*) = *F_A_*(*γ*(*θ*)) – *γ*(*f_A_*(*θ*)).
2. Compute *∂_θγ_*(*f_A_*(*θ*)) = *TX*(*γ*(*f_A_*(*θ*))).
3. Compute 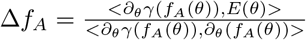.
4. Set *f_A_*(*θ*) ← *f_A_*(*θ*) + Δ*f_A_*(*θ*).
5. Repeat steps 1-4 until the error *E* is smaller than the established tolerance. Then *PRC*(*A, θ*) = *f_A_*(*θ*) – (*θ* + *T_rel_*/*T*).

We refer the reader to [50] for the implementation of this methodology for not pulsatile perturbations. To compute the iPRC by means of this algorithm one has to consider perturbations of *A* small and *f_A_*(*θ*) = *θ* + *T_rel_*/*T* as initial seed. Then, ∇Θ(*γ*(*θ*)) = *PRC*(*A, θ*)/*A*.

## Acknowledgements

AP and JH acknowledge the long-term strategic development financing of the Institute of Computer Science (RVO:67985807) of the Czech Academy of Sciences, JH acknowledges the support of project Nr. LO1611 with a financial support from the MEYS under the NPU I program. The funders had no role in study design, data collection and analysis, decision to publish, or preparation of the manuscript.

